# RevGel-seq: instrument-free single-cell RNA sequencing using a reversible hydrogel for cell-specific barcoding

**DOI:** 10.1101/2022.07.01.498266

**Authors:** Jun Komatsu, Alba Cico, Raya Poncin, Maël Le Bohec, Jörg Morf, Stanislav Lipin, Antoine Graindorge, Hélène Eckert, Azadeh Saffarian, Léa Cathaly, Frédéric Guérin, Sara Majello, Damien Ulveling, Anaïs Vayaboury, Nicolas Fernandez, Dilyana Dimitrova, Xavier Bussell, Yannick Fourne, Pierre Chaumat, Barbara André, Elodie Baldivia, Ulysse Godet, Mathieu Guinin, Vivien Moretto, Joy Ismail, Olivier Caille, Natacha Roblot, Carine Beaupère, Alexandrine Liboz, Ghislaine Guillemain, Bertrand Blondeau, Pierre Walrafen, Stuart Edelstein

## Abstract

Progress in sample preparation for scRNA-seq is reported based on RevGel™-seq, a reversible-hydrogel technology optimized for samples of fresh cells. Complexes of one cell paired with one barcoded bead are stabilized by a chemical linker and dispersed in a hydrogel in the liquid state. Upon gelation on ice the complexes are immobilized and physically separated without requiring nanowells or droplets. Cell lysis is triggered by detergent diffusion, and RNA molecules are captured on the adjacent barcoded beads for further processing with reverse transcription and preparation for cDNA sequencing. As a proof of concept, analysis of PBMC using RevGel-seq achieves results similar to microfluidic-based technologies when using the same original sample and the same data analysis software. In addition, a clinically relevant application of RevGel-seq is presented for pancreatic islet cells. Furthermore, characterizations carried out on cardiomyocytes demonstrate that the hydrogel technology readily accommodates very large cells. Standard analyses are in the 10,000-input cell range with the current gelation device, in order to satisfy common requirements for single-cell research. A convenient stopping point after two hours has been established by freezing at the cell lysis step, with full preservation of gene expression profiles. Overall, our results show that RevGel-seq represents an accessible and efficient instrument-free alternative, enabling flexibility in terms of experimental design and timing of sample processing, while providing broad coverage of cell types.

---

Research in many areas of biology focuses on single-cell studies, particularly single-cell RNA sequencing (scRNA-seq) mainly using microfluidic-based methods^1–6^, with the 10x Genomics Chromium instruments achieving particularly wide usage^7^. Comparative benchmarking has been reported for many of the available methods (including CEL-Seq2, MARS-Seq, Quartz-Seq2, gmcSCRB-seq, Smart-Seq2, C1HT-small, C1HT-medium, Chromium, ddSEQ, Drop-Seq, ICELL8, and inDrop)^8–10^. Major efforts using scRNA-seq have been applied to define atlases, notably for human and mouse cells^11, 12^, with a potential for assessing cell-cell and virus-host interactions^13,14^, facilitated by technical optimization of methodologies^8,15^. Other investigations have focused on specific organs, often in relation to pathologies^16^, especially cancer, immunotherapy, and cardiovascular diseases^17–19^. Research has focused as well on development ^20,21^, various forms of multiomics^22^, and spatial transcriptomics^23^. In addition, applications to plants are emerging^24^, highlighting single-nucleus (snRNA-seq) approaches^25^.

While these applications have produced important new findings, further optimization of scRNA-seq sample preparation could provide the following benefits:

1. Immediate processing of fresh samples with transcriptomes unaltered by freezing, fixation, or mechanical stress, while permitting instrument-free sample preparation on multiple sites for clinical and collaborative studies or in biohazard laboratories;
2. Enabling convenient stopping points to permit interruption of processing for later analysis, while fully maintaining transcriptome integrity;
3. Providing scalability for different numbers of cells in a sample;
4. Providing compatibility with cells of different sizes, especially very large cells;
5. Monitoring of initial cell and barcoded bead complex formation to assure sufficient yield to proceed, with the possibility of initially sequencing a fraction of the sample for quality control purposes.

RevGel-seq is a scRNA-seq preparation method based on a reversible hydrogel technology that provides advances on most of the above points. The workflow, presented in Fig. 1A, comprises liquid-gel-liquid transitions of the reversible hydrogel that contains cells linked to barcoded beads. Sequencing outputs are evaluated and displayed using Cytonaut (https://cytonaut-scipio.bio), a cloud-based data analysis platform that includes pre- and post-processing, as well as interactive data visualization (see Fig. 1B). Being a cloud solution, Cytonaut can be used via a simple web interface, bypassing the need to install software packages or use computational resources. Overall, results presented establish RevGel-seq performance levels, in comparison to the predominant microfluidics technology, and illustrate a clinically relevant application.

**Figure 1.**
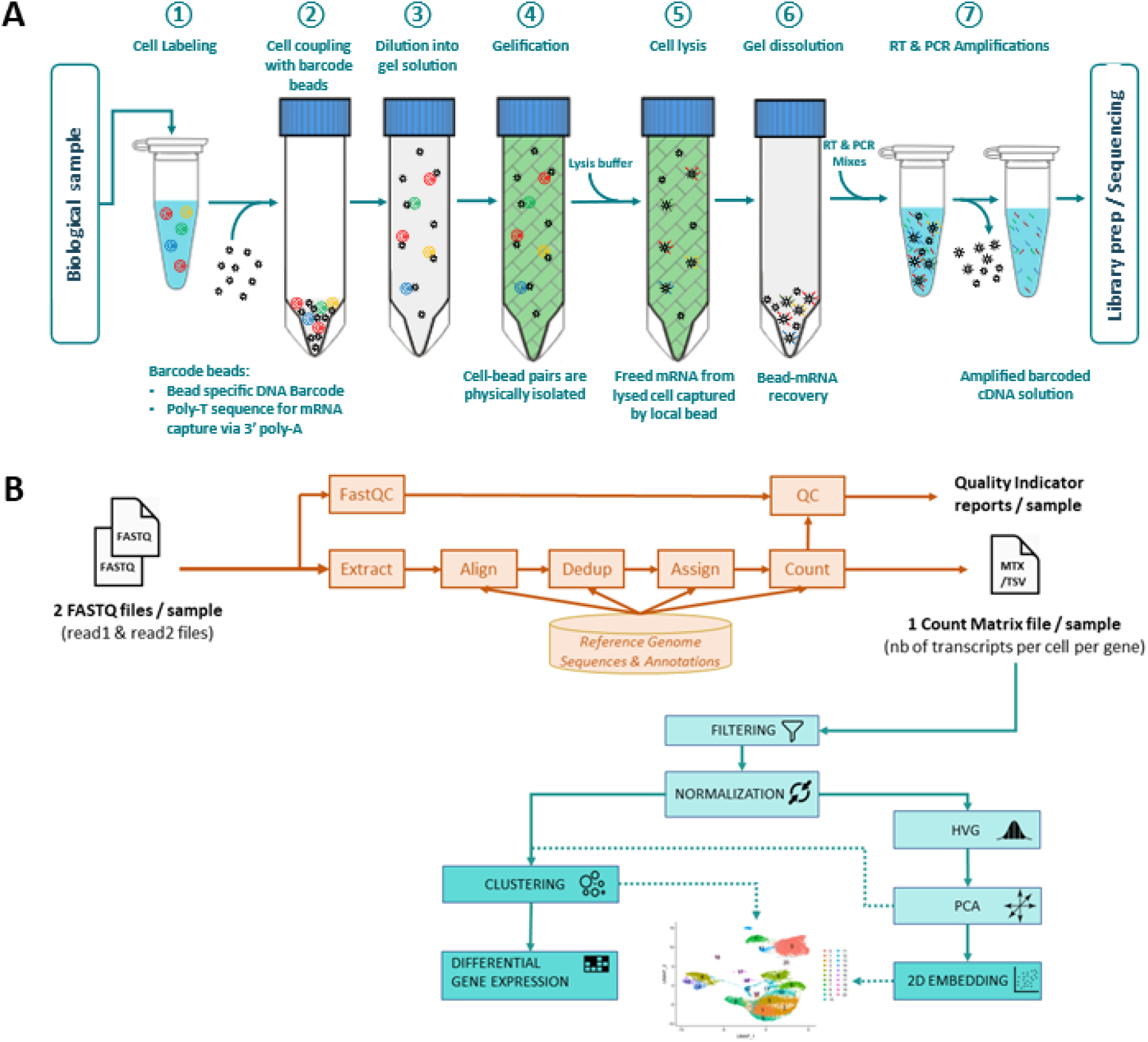
Sample preparation workflow and data analysis pipeline. (**A**) RevGel-seq workflow steps for sample preparation to characterize scRNA-seq. Barcoded beads and cells are attached via a bifunctional chemical linker. These complexes are dispersed in the hydrogel in the liquid state and immobilized upon gelation. Following cell lysis, RNA molecules are captured on the barcodes. After reverse transcription, barcoded cDNAs are PCR-amplified. Further details are presented in Methods. (**B**) Workflow of the end-to-end data processing pipeline integrated in the Cytonaut platform. The pre-processing phase inputs the raw sequencing data (FASTQ files) and outputs quality indicators and count matrices, followed by the post-processing phase that inputs the count matrices to perform 2D embedding, cell clustering and differential gene expression. The Cytonaut Rover module enables interactive data visualization. Additional details in Methods.

## Results

### Mixed species experiment

The performance of RevGel-seq was assessed using a cell barcode purity criterion (see Methods) by Cytonaut (v1.2) in the sequencing data for each of the three technical replicates. Fig. 2A represents the Barnyard plot for one of the technical replicates, where cell barcodes are annotated based on the percentage of detected transcripts from each species. The “human” cells (in orange) and “mouse” cells (in teal) are cell barcodes for which cell barcode purity is higher than 95%. The transcripts from the two types of cells were readily separated, with a hetero-species cell multiplet rate of 3%, implying an equal rate of non-detectable homo-species cell multiplets. A hetero-species cell multiplet is defined as a barcode for cell barcode purity is lower than 2/3.

**Figure 2.**
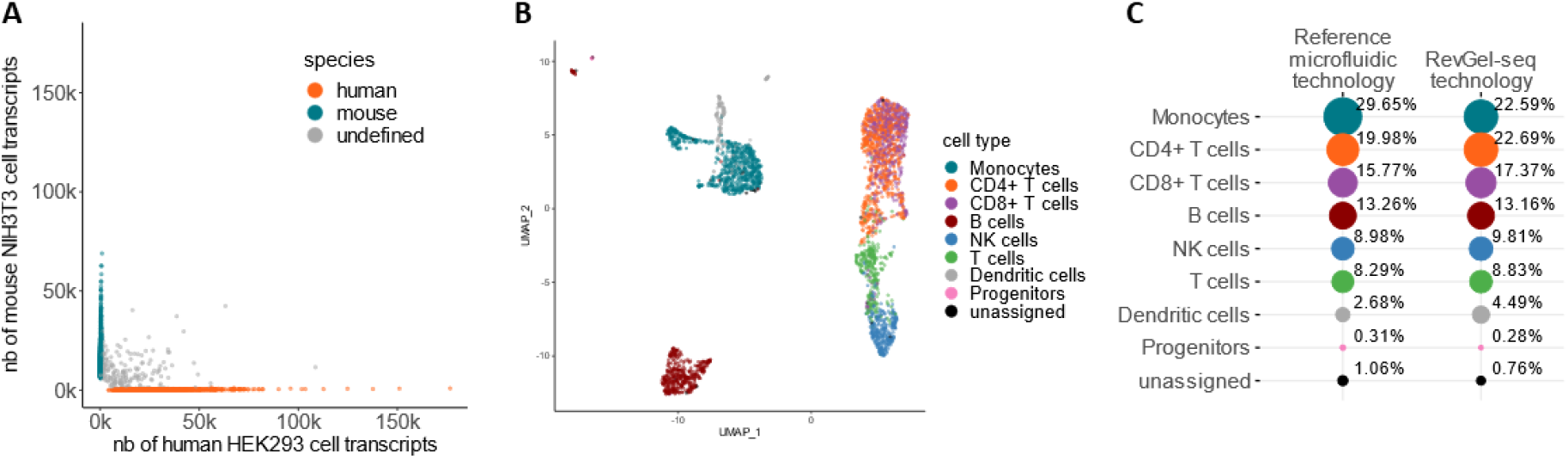
Benchmarking and applications. (**A**) Barnyard plot showing for each cell-associated barcode the number of detected mouse NIH3T3 transcripts and the number of detected human HEK293 transcripts, from 10,000 input cells prepared with RevGel-seq, with sequencing data downsampled by raw read subsampling at a depth of 50,000 raw reads per cell on average. The hetero-species cell multiplet rate is 3% (for details on criteria see Methods). (**B**) Cell classification and 2D embedding from a PBMC sample of 10,000 input cells prepared with RevGel-seq were downsampled by raw read subsampling at a depth of 20,000 raw reads per cell on average. Post-processing was performed using Seurat^21^ and automated cell classification was performed using the SingleR^22^ algorithm based on the reference dataset MonacoImmuneData^23^. Unassigned cells had classification uncertainties that were considered too high according to the pruneScores method with default parameters. All relevant PBMC sub-types were identified. (**C**) Percentage of automatically classified cells for each cell type identified in the same PBMC sample prepared with a reference microfluidic technology, (10x Chromium 3’ v3.1) and prepared with RevGel-seq, with 10,000 input cells and a sequencing depth of 20,000 raw reads per cell on average. For both methods, RevGel-seq and 10x Chromium 3’ v3.1, the same original sample was used and the same data analysis software (Cytonaut v1.2) was applied. The relative proportions of cell types are highly similar between RevGel-seq and the reference microfluidic technology.

### Performance comparison with 10x Chromium 3’ v3.1 on human PBMC

To evaluate RevGel-seq on samples with multiple cell types, analyses were performed on peripheral blood mononuclear cells (PBMCs)^26^. From the same anonymized healthy blood donor (see Methods), input cells of human PBMCs were used for the RevGel-seq procedure (10,000 input cells) and for the 10x Chromium with the single-cell 3’ reagent kit (v3.1) procedure (10,000 input cells). For both sample preparation methods, the same data analysis pipeline (Cytonaut v1.2) was applied and sequencing data were downsampled to ~20,000 raw reads per cell on average, as described in the Methods section. Cell type annotation was performed with SingleR (v1.8.1)^27^ using MonacoImmuneData^28^ as a reference of cell types and gene expression. The results are presented as a UMAP 2D projection with color-coded cell types in Fig. 2B and as annotated cell type proportions in Fig. 2C. The automated cell-wise classification demonstrates that RevGel-seq and a 10x Chromium 3’ v3.1 both identified the same cell types, with the same hierarchy of abundance. Some differences in the percentage of the population for several cell types were observed among a total of 3,160 analyzed cells for the RevGel-seq sample and 4,509 analyzed cells for the 10x sample. Expression profiles of selected marker genes of PBMC show good concordance between the results obtained with RevGel-seq and 10x Chromium 3’ v3.1 (Fig. S1).

### Characterization of pancreatic islet cells

An application of RevGel-seq to a current pre-clinical research project is illustrated by studies to phenotype pancreatic islet cells in relation to insulin resistance and islet cell adaptation^29^. After sequencing data analysis with Cytonaut (v1.2), the automated classification of pancreatic cell types (Fig. 3A) and associated gene counts (Fig. 3B) are in accord with previous studies^30^. Expression profiles of known marker genes^30^ are presented in Fig S2, revealing the predominant gene expression for each cell-type, notably for the three principal cell-types alpha, beta, and delta: glucagon (GCG), insulin (INS), somatostatin (SST), respectively.

**Figure 3.**
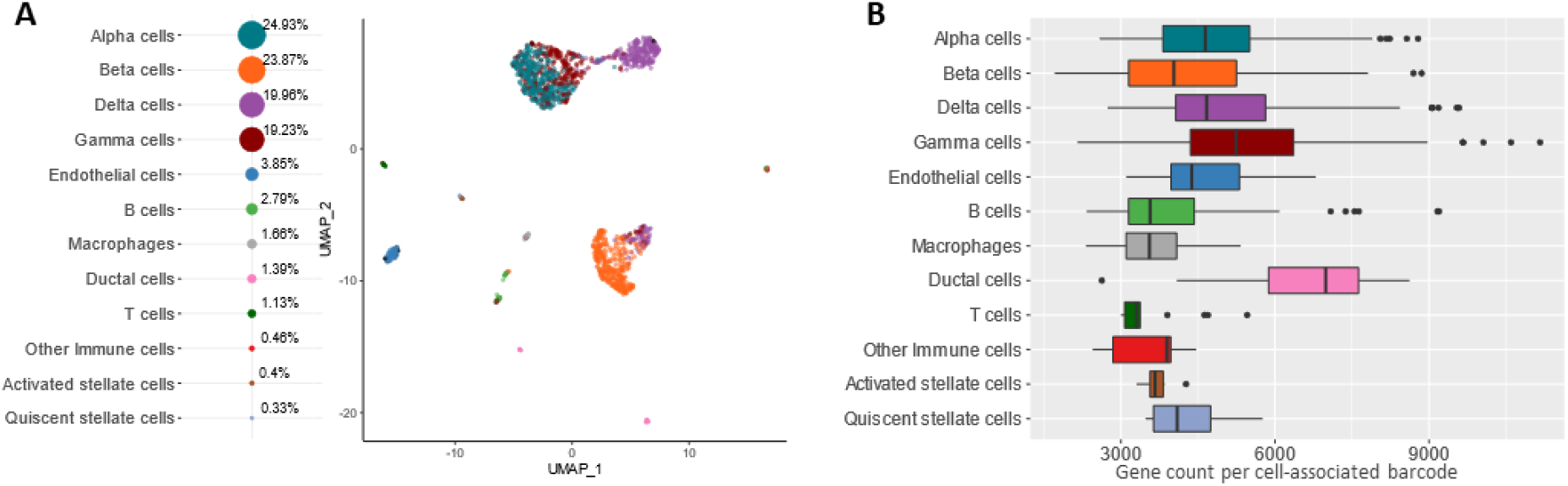
Analysis of pancreatic islet cells. (**A**) Pancreatic islet cells (176,000 raw reads per cell) automatically classified into cell types (left) following the same methodology as in Fig. 2B except for the reference dataset BaronPancreasData^30^. (**B**) Box plots showing the distribution of gene diversity per identified cell type for pancreatic islet cells classified in Fig. 3A.

### Establishment of large cell compatibility

We next tested RevGel-seq compatibility with large cells, using cardiomyocyte suspensions. Bead-cell complexes were subsampled and examined under brightfield microscope. A cardiomyocyte cell suspension prior to coupling to barcoded beads is illustrated in Fig. 4A, with the complexes of cardiomyocytes and barcoded beads after coupling presented in Fig. 4B. Counting coupled cell-bead objects in this subsample provided an estimate of the quantity processed to cDNA. The numbers of barcoded beads in the eight final PCR tubes (see Methods, section 7v) were also counted to estimate the bead recovery rate. The product of the number of cell-coupled barcoded beads and the bead recovery rate provided an estimated number of cells to be found in the corresponding sequence data. Since the protocol separates the barcoded beads sample into eight PCR tubes, which is equivalent to splitting the input cells into eight sub-fractions, either all the input cells or only a fraction can be sequenced, depending on needs or for Quality Control purposes before sequencing the entire sample. For the current study, barcoded cDNAs from two PCR tubes were converted to the sequencing library separately (each represents 1,250 input cells). For both subsamples #1 and #2, the number of cells found in the output data (530 and 523, respectively) was consistent with the number of bead-cell complexes estimated by microscope observations (595 and 527, respectively), as shown in Fig. 4F, demonstrating that microscopic assessment of cell-barcoded bead complexes enables a useful estimation of the number of cells in sequencing data.

**Figure 4.**
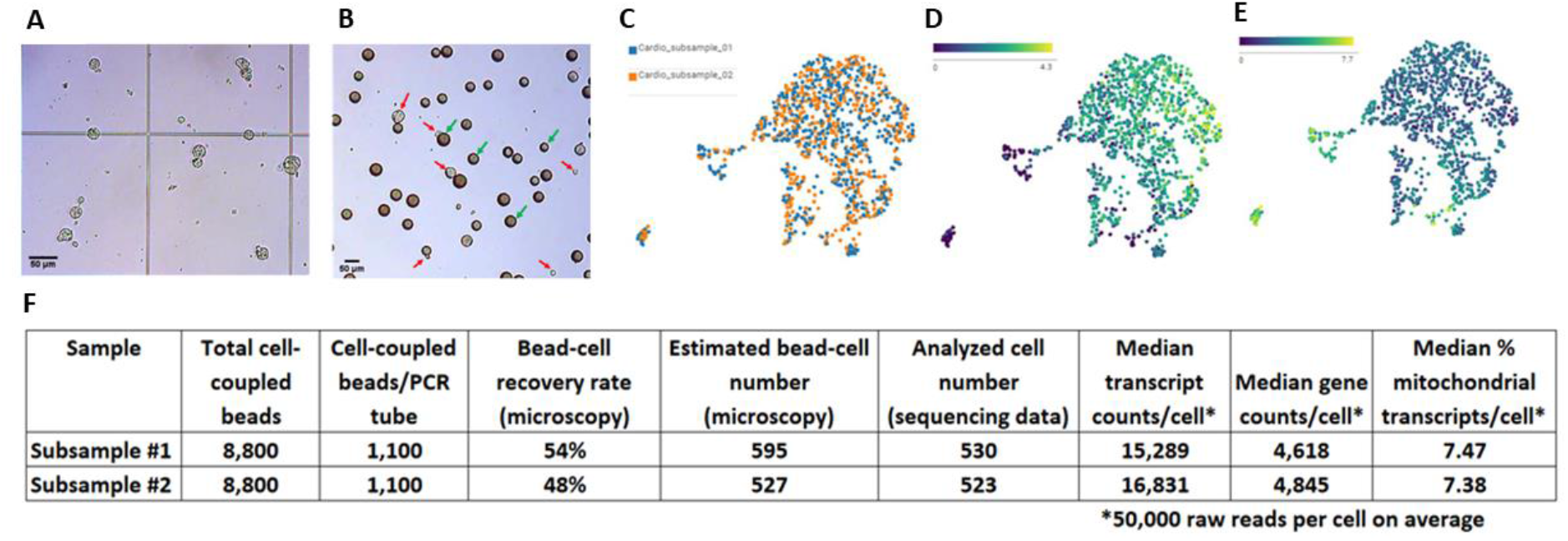
Evaluation of large cell processing with cardiomyocytes. (**A**) Microscopic observations of cardiomyocytes following trypsinization (cell diameter of 20-35 μm). (**B**) Microscopic observations of cardiomyocytes following coupling procedure (red arrows indicate cells, green arrows indicate beads). (**C**) 2D projections of data from two subsamples (PCR tubes) from the same cardiomyocyte sample preparation. (D) Gene expression of RYR2 (cardiomyocyte enriched gene) and (E) CALD1 gene expression (smooth muscle cell enriched gene). (**F**) Estimated cell quantities obtained from numbers of cell-bead complexes (for all 10,000 input cells), observed bead recovery rate (for each PCR tube, max 12.5% for each of 8 PCR tubes), and transcript/gene counts per analyzed cell at average 50,000 raw reads per cell. Sequencing data was processed according to Section I above by the pre-processing analysis pipeline of Cytonaut which automatically detects the analyzed cells.

The raw sequencing data of the two subsamples were analyzed by Cytonaut (v1.2) which performed initial data pre-processing to generate count matrices and quality indicators, then data post-processing with dimension reduction and projection in 2D. The UMAP graphs presented in Fig. 4C show that cells from both subsamples are homogeneously distributed, indicating the absence of batch effects of split cDNA amplification and subsequent library preparation. Results in Fig. 4F demonstrate that the median number of transcripts, the median number of genes, and the median mitochondrial transcript rate per cell are all comparable between the two libraries. The gene expression levels of RYR2 and CALD1 genes are presented in Fig. 4D and Fig. 4E respectively. The levels of expression of the two genes are well contrasted, implying that most cells are cardiomyocytes (RYR2 high; Fig.4D), with a minor fraction of smooth muscle cells (CALD1 high; Fig.4E).

### Determination of early stopping point

To establish the feasibility of pausing the sample preparation at an early stage (2 hours), a comparison of two samples frozen in the gelation device (one, directly at −80°C freezer, and the other, flash freezing in dry ice) and an unfrozen control sample was performed by sequencing of one cDNA PCR tube (equivalent to 1,250 input cells) from each of three samples, at the point in the protocol indicated in Fig. 5A. All representative fundamental performance indicators computed with Cytonaut v1.2 are comparable between the unfrozen control sample and the two frozen samples, as indicated in Fig. 5B. The count matrix tables of all three samples were merged then subjected to PCA and 2D projection on UMAP in order to see whether cells from samples stopped at 2 hours are distributed similarly compared to control. Results in Fig. 5C show that cells from all three different conditions are uniformly distributed on the UMAP, while keeping good segregation of human and mouse cell identity as presented in Fig. 5D. These analyses confirm that freezing of cell-bead complexes in the hydrogel at −80°C overnight did not alter the cell capture efficiency nor the transcriptome integrity.

**Figure 5.**
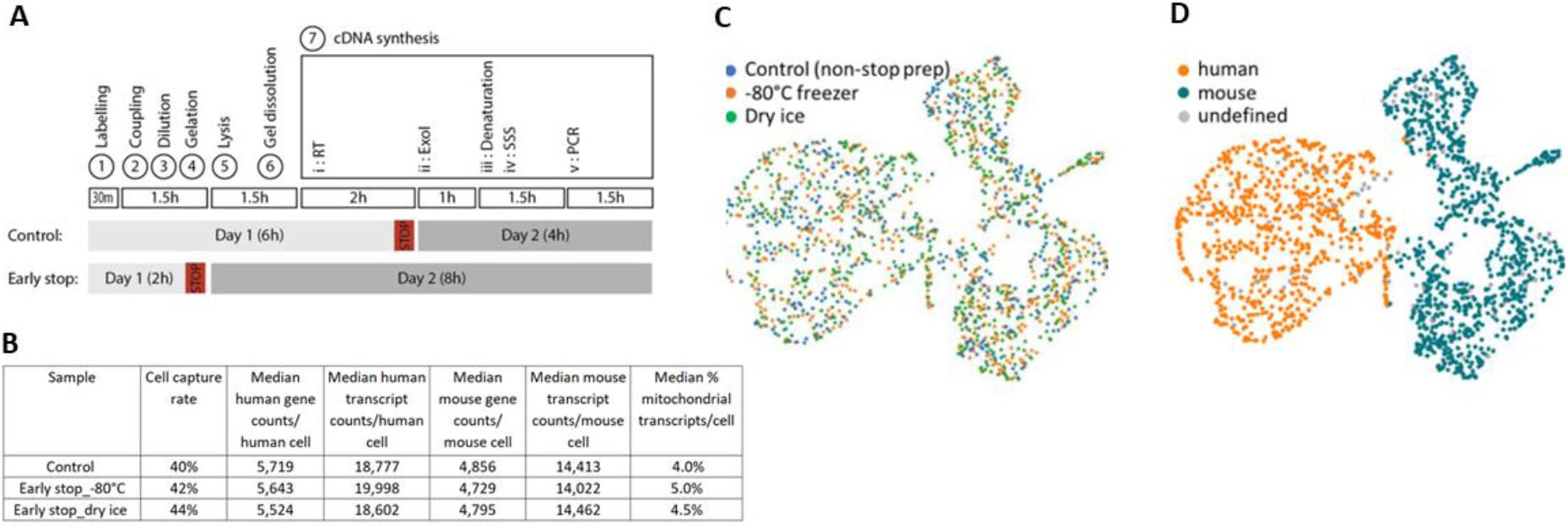
Freezing samples at an early stopping point during RevGel-seq protocol does not modify scRNA preparation performances. (**A**) Schematic representation of the timing differences between the “Control” and “Early stop” samples (See Methods for specific steps). “Control” samples were obtained by freezing the RT reaction mixture at −20°C overnight, while “Early stop 80°C” and “Early stop dry ice” samples were obtained by freezing (either at −80°C or in dry ice, respectively) immediately after applying lysis buffer to the hydrogel. The RevGel-seq protocol was resumed on a subsequent day for all samples. (**B**) Comparison of performance indicators between the tested conditions (average of 3 PCR tubes for each condition). Gene and transcript counts per analyzed cell at 50,000 raw reads per cell with raw read downsampling. (**C, D**) Projection of scRNA-seq data on 2D (UMAP) of the pooled samples (pool of data from 1 PCR tube for all 3 conditions), shown per condition (**C**) and per determined cell species (**D**). (“undefined” corresponds to cells with a cell barcode purity lower than 95%; see Section VIII A-iii of data processing above). With respect to cell capture yields, obtained cell species proportions were as follows: human 45.8%, mouse 49.9%, undefined 4.3% (Control sample); human 48.6%, mouse 52.5%, undefined 3.9% (Early stop dry ice); and human 46.2%, mouse 48.4%, undefined 5.4% (Early stop −80°C).

## Discussion

In this study, we present results obtained using the RevGel-seq approach, a single-cell RNA instrument-free sample preparation technology that relies on formation of cell-barcoded bead complexes. The complexes are suspended in a gelation tube containing a reversible hydrogel in the liquid state. Upon transition to the gel state by incubating on ice, the complexes are separated by the gel matrix. For typical conditions of experiments presented here (for 10,000 cells in 2 mL of gel) the average center-to-center distance between cells is ~500 microns, sufficient to minimize crosstalk (mRNA from one complex diffusing after cell lysis to the barcoded bead of another complex in the vicinity). Following cell lysis and degelation, barcoded beads are recovered and the information combining mRNA 3’ end and barcode sequence is obtained through a series of biomolecular steps (reverse transcription, ExoI digestion, second strand synthesis and PCR amplification), leading to single cell cDNAs ready for library preparation and subsequent sequencing.

RevGel-seq performances have been assayed using mixed-species cell suspensions, human peripheral blood mononuclear cells, and pancreatic islet cells. Mixed species RevGel-seq experiments showed an efficient segregation between human and mouse cells with a hetero-species cell multiplet rate estimated at 3% (Fig. 2A), while RevGel-seq assays using PBMCs (Fig. 2B) and pancreatic islet cells (Fig.3) identified marker gene expressions and cell type proportions comparable to those obtained using 10x Chromium 3’ v3.1 and to previously published results^30^, respectively, although with lower capture rates by RevGel-seq for the former. Nevertheless, our results show that the RevGel-seq approach is a convenient and efficient sample preparation alternative for scRNA-seq studies.

Within the four areas of potential improvements noted in the introduction, the RevGel-seq technology provides benefits that can facilitate scRNA-seq studies for specific applications.

First, RevGel-seq demonstrated successful sample preparation for scRNA-seq without any specific instrument. Since only standard laboratory equipment is required, RevGel-seq sample preparation can be carried out at any sample collection site. This feature eliminates not only an upfront investment to dedicated equipment but also any need of fixation or freezing of cells, as well as avoiding mechanical stress (all of which are known to alter gene expression profiles), and processing delays caused by the transfer of samples to another facility/laboratory.

Second, a 2-hour stopping point (Fig.5), offers the possibilities of preparing samples at any time of need and centralizing final processing for samples collected at different times or different locations.

Third, concerning scalability, the number of cells that can be analyzed with RevGel-seq is dependent on the current gelation device design. Although scalability can be achieved by other methodologies (SPLiT-Seq, PIP-seq), the current level of cell processing for RevGel-seq is not a conceptual limitation. Increase of hydrogel volume is readily possible by modifying the gelation device to multiply the number of cells processed at once.

Fourth, compatibility with cells of any size is inherently achieved, since separation of cell-bead complexes by the surrounding hydrogel enables processing of large cells, as demonstrated for cardiomyocytes (Fig.4). In contrast, microfluidic-or nanowell-based methods are limited in their ability to process large cells.

Fifth, the captured beads can be subsampled following the PCR amplification step: the option of sequencing a subfraction of the sample (i.e., an individual PCR tube instead of the pool of PCR tubes) provides the opportunity to obtain solid preliminary results at limited cost. As shown by the inter-tube repeatability results, good quality control results of sample fraction sequencing ensure the quality of whole sample sequencing at same or higher sequencing depth.

In conclusion, RevGel-seq achieves flexibility in scRNA sample preparation by circumventing difficulties that researchers currently experience, notably eliminating the need for special instruments, allowing sample preparation without constraints on timing and location, permitting subsample analysis to optimize sequencing costs, and accommodating large cells for processing. Dedicated benchmarking experiments will be needed to comprehensively compare RevGel-seq to the other recently developed technologies^31–33^ in terms of performance (cell capture and mRNA capture efficiency; cell multiplet rate; range of cell types analyzable), accessibility (availability at time of need, utilization in normal and restricted environments) and convenience (protocol complexity, total processing time).

## Methods

### I. RevGel-seq sample preparation workflow

Experiments were performed with the RevGel-seq protocol, capable of analyzing 10,000 input cells per sample with the specially designed gelation device (Fig. S3). The individual steps shown in Fig. 1A, from cell-barcoded bead coupling to library preparation for sequencing, are more fully described below:

#### 1) Cell labeling

Cells are first labelled with a bifunctional chemical linker used to tether individual cells to barcoded beads. The cells are incubated with this bifunctional linker (polyA at one extremity and a hydrophobic moiety at the other extremity) in DPBS for 5 min. The labelled cells are washed by 0.1% BSA in DPBS twice and resuspended at a density of 100 labelled cells/μL with the same buffer for a sample preparation with 10,000 input cells.

#### 2) Cell coupling

For preparation of the cell-bead complexes, 100 μL of the barcoded bead suspension^1^ (1,000 beads/μL, 20% PEG8000 in DPBS) are pipetted into the bottom of gelation tube (see Fig. S3), followed by transfer of 100 μL of the labelled cell suspension into the suspension of barcoded beads. The combined suspension is homogenized by micropipette to couple the individual labelled cells to individual barcoded beads upon their collisions. The number of barcoded beads is in large excess to the number of labelled cells to avoid that one bead is bound to more than one cell. Quantification of cell-bead complexes can be achieved at this point by microscopy, if evaluation of coupling is required.

#### 3) Dilution into gel solution

Cell-bead complexes suspension (200 μL) are diluted into 10x volume of hydrogel solution (200 μL). The hydrogel solution contains thermal sensitive polymers and is isotonic with a density matched to barcoded beads to prevent cells from osmotic shock as well as beads from precipitation prior to gelation. The diluted cell bead complexes are then homogenized by simple rotation of the gelation tube. After this homogenization, the gelation piston is slowly inserted into the gelation tube while keeping both vertical, as the contents rise smoothly along the wall of the gelation tube with a level horizontal meniscus.

#### 4) Gelation

The assembled gelation device is then placed vertically on ice and incubated for 20 min for complete gelation to immobilize the cell-bead complexes. This resulting thin hydrogel layer has a high surface to volume ratio to make the subsequent cell lysis efficient and synchronous for all cells. While maintaining verticality, the gelation piston is slowly removed from the gelation tube. The gel remains in the tube, partially collapsed, with no gel remaining on the piston. Then 7.5 mL of lysis buffer (sarkosyl 0.2% w/v, 50 mM dithiothreitol, 20 mM EDTA and 200 mM Tris pH 7.5) are added to the gelation tube. If necessary, the procedure can be interrupted at this two-hour stopping point by placing the gelation device vertically in a freezer at −80°C or flash freezing in dry ice, to resume the experiment at a later time by thawing the frozen device at 25°C water bath for 15 min.

#### 5) Cell lysis and mRNA capture

The hydrogel sheet in the lysis buffer is incubated at room temperature for 1 hour with a gentle orbital shaking. Cell lysis reagents diffuse into the hydrogel and lyse cells in the gel, triggering release of polyA tailed RNA and their hybridization on the 3’-polyT extremities of the barcoding oligonucleotides on the capture bead coupled to each cell. The released mRNA molecules are confined in the vicinity of each cell by the hydrogel polymer mesh.

#### 6) Gel dissolution

When lysis is completed, the degelation step is immediately initiated by adding 3.75 mL of degelation buffer (3M guanidine thiocyanate and anti-foam reagent supplied to the reverse transcription reaction buffer (RT buffer)) in the gelation tube and sealed with the gelation tube cap. The gelation tube is then held in the vertical position and vortexed for 5 min at maximum speed. The barcoded beads with hybridized mRNA are now free from hydrogel and centrifuged at 1,000 x g for 3 min.

#### 7) RT, PCR, and preparation for DNA sequencing

The RNA-loaded barcoded beads are washed once with 5 ml of ice-cold RT buffer, and transferred into 1.5 mL microtubes, then washed two additional times to prepare for subsequent enzymatic steps.

i. Reverse transcription: The pellet of beads is resuspended in 20 μL of bead wash buffer and supplemented by 75 μL of the RT supermix composed of 1.3 x RT buffer supplied with Maxima H Minus reverse transcriptase (ThermoFisher, EP0751), 1.3 mM each dNTP, with 100 U of NxGen RNase inhibitor (LGC Biosearch technologies, 30281), 2 U of beta-agarose I (NEB, M0392), and 5 μL of Maxima H minus reverse transcriptase. The sample is then placed in a heat block and incubated 10 min at 25°C followed by 90 min at 42°C. Next, the sample tube is placed on ice for 2 min and then spun briefly on a benchtop centrifuge. The procedure can be interrupted at this point and the sample stored overnight at 4°C or up to one week at −20°C. When ready to continue, the sample is thawed on ice before proceeding to the next steps.
ii. Endonuclease I treatment: 5 μL of Exonuclease I is added to the sample and homogenized by pipetting. The remaining capture oligos non-hybridized with captured RNA are digested by incubation for 50 min at 37°C. After this incubation the enzymes are removed by washing with 250 μL of TE (10mM Tris, 1mM EDTA, pH 8) with 0.5% (w/v) SDS.
iii. Alkaline denaturation: The beads are washed once with 100 μL 0.1M NaOH solution and resuspended in 50 μL 0.1M NaOH solution. RNA molecules are then removed from cDNA by 5 min incubation on a microtube rotator/wheel. The bead washing is then carried out with 250 μL of TE-TW (10mM Tris, 1mM EDTA, pH 8; 0.01% Tween 20) and adjusting the residual volume to 20 μL using RT buffer.
iv. Second strand synthesis: The S3 supermix (1.1x RT buffer, 1.3 mM each dNTP, 12% PEG8000, 13 μM second strand primer) is first incubated for 5 min at 70°C, then immediately placed on ice for 1 min, spun, and homogenized. Then 77.5 μL of the S3 supermix is added to the bead suspension, followed by addition of 2.5 μL of the S3 enzyme. The sample is thoroughly homogenized by pipetting and incubated 1 hour at 37°C. Two washes with 250 μL of TE-TW are carried out, followed by a final resuspension in 160 μL with nuclease-free water. Relevant oligonucleotides are described in Table S1.
v. Polymerase chain reaction: 240 μL of the PCR supermix (1.66x KAPA HiFi HotStart readymix (Roche, 07958935001), 1.33 μM PCR primer) is added to the 160 μL of the bead suspension. The beads in the reaction mix are homogenized by pipetting and split into 8 PCR tubes, with 50 μL in each tube. The PCR program is described in Table S2.
vi. PCR strip tubes are spun for 20 sec using a benchtop centrifuge, and 45 μL of bead-free supernatant from each tube is transferred to another clean PCR tube separately. Size selective purification is performed at 0.6x volume of SPRIselect reagent (Beckman Coulter, B23317) to the transferred PCR product (27 μL SPRIselect to 45 μL of the PCR product) and following the manufacturer’s procedure. Elution is performed in 20-40 μL of nuclease-free water. Quality check of the purified cDNA amplicons is made by Agilent’s TapeStation HSD5000. The purified cDNA can be stored at −20°C, then thawed on ice before proceeding to the next steps.
vii. Library preparation: 600 pg of the purified cDNA amplicons from each PCR tube are sampled and processed according to Illumina Nextera XT library preparation kit (illumina, FC-131-1024) using a custom library prep primer for the P5/read1 side of the library amplification PCR.

### II. Mixed species sample preparation

Both human cell line HEK293 and mouse cell line NIH3T3 were cultured separately in complete DMEM media. Both cell lines were harvested, and the cell density was adjusted to 100 cells/μL. Each cell suspension was mixed at 1:1 ratio to obtain mixed cell suspension at 50 human cells and 50 mouse cells/μL. This mixed cell suspension was used as start material of RevGel-seq sample prep for species mix experiment.

### III. PBMC sample preparation

Whole human blood from an anonymous healthy donor was provided from the *Etablissement Français du Sang (EFS)* under a convention for research use only. PBMCs were isolated from fresh blood by HistoPaque-1077 (Sigma) according to the instructions. The purified PBMC from the same blood sample was used for both the RevGel-seq method described above (with doubling of the ratio of beads per cell to increase coupling yield), and the 10x Genomics Chromium single cell 3’ reagent kit (v3.1), and both scRNA sample preparations with 10,000 input cells were started at the same time.

### IV. Pancreatic islet cells

Pancreas islets were isolated from mouse (8 weeks male C57BL/6J) pancreas by collagenase (1 mg/ml, Sigma-Aldrich) injection in the bile duct and handpicked under a binocular microscope (Leica). Islet cells were dissociated with 0,05% trypsin-EDTA (Gibco Thermo Fisher), incubated at 37°C for 5 min and resuspended in PBS with 2% fetal calf serum and 0,5 mM EDTA. Two technical replicates of 10,000 input cells from this pancreas islet were prepared following RevGel-seq method described above.

### V. Cardiomyocytes, including an early checkpoint

Human cardiomyocyte cells were purchased from Takara Bio (ref Y10060). The cells were cultured according to the instructions and harvested by trypsin just before processing with the RevGel-seq method described above with 10,000 input cells. After the cell-bead coupling step, 20 μL of the cell-bead complex suspension was sampled with a wide-bore pipette tip and transferred to a cell counting chamber for microscopic examination of the cell-bead complex state. The remaining cell bead complex suspension was processed according to the RevGel-seq procedure and sequenced.

### VI. Early stopping point

To confirm the possibility to safely stop the experiment at an early stopping point in the RevGel-seq procedure, three samples of 10,000 input cells of HEK293 and NIH3T3 in a 1:1 ratio were prepared from the same mixed cell suspension and their preparation was stopped at two different times. One sample was stopped after reverse transcription as a known stopping point^1^. The two other samples were stopped immediately after applying lysis buffer to the hydrogel (2 hours from the start of sample preparation) by freezing, one in a −80°C freezer, and the other on dry ice for 20 min then transferred to −80°C freezer. The following day, the two frozen samples in the cell lysis buffer were thawed at 25°C in water bath for 15 min prior to resuming the remaining sample preparation procedure.

### VII. Sequencing

All sequencing of RevGel-seq samples was performed on a NovaSeq 6000 with spiking custom primer for read1. The read length was 30 bases for read1, 8 bases for i7 index, and 70 bases for read2. Sequencing of 10x Chromium 3’ RNA samples was performed according to the instructions and NovaSeq 6000 was used as well.

### VIII. Data analysis with Cytonaut, the Scipio bioscience cloud-based bioinformatics platform

The end-to-end data analysis from raw sequencing data to differential gene expression is performed by Cytonaut (v1.2) cloud software (www.cytonaut-scipio.bio), which integrates data pre-processing (v6.7) as well as data post-processing (v1.5) and interactive data visualization (v1.2). The same end-to-end data analysis methodology based on Cytonaut was applied to all samples except for RevGel-seq and 10x PBMCs samples and for RevGel-seq Pancreatic islet cells samples for which Cytonaut was used for pre-processing analysis but a separate script was used for post-processing analysis based on Seurat (v4.1.0) and automated cell annotation based on SingleR (v1.8.1). The workflow of the end-to-end data processing pipeline integrated in the Cytonaut platform is summarized in Fig. 1.b, with details on individual steps presented below:

#### A) Pre-processing pipeline

The data pre-processing pipeline of Cytonaut takes as input the two FASTQ files of the sample (R1 and R2), leverages the specific barcode pattern of RevGel-seq capture beads, and successively performs read quality control, detection of cell-associated barcodes, read alignment based on exons only, read deduplication, and read assignment. Detection of cell-associated barcodes was performed for each sample by applying the distance-based knee method of UMI-tools v1.1.2. The reference genome sequences and annotations are based on Ensembl release v99 for human species (GRCh38; ftp.ensembl.org//pub/release-99/fasta/homo_sapiens/dna/; ftp.ensembl.org//pub/release-99/gtf/homo_sapiens/) and mouse species (GRCm382;https://ftp.ensembl.org/pub/release-99/fasta/mus_musculus/dna/; ftp.ensembl.org//pub/release-99/gtf/mus_musculus/).

The pre-processing pipeline provides as output:

i. The count matrix of the input sample, which contains the number of transcripts detected for each gene in each detected cell;
ii. A set of quality indicators, which include in particular the median number of genes per cell, the median number of transcripts per cell, the median mitochondrial transcript rate per cell, as well as their distribution statistics;
iii. For the NIH3T3-HEK293 sample in Fig. 2.a, the quality indicators also include statistics on cell barcode purity. Cell barcode purity is the probability that a transcript captured by the cell barcode has been expressed by the main cell coupled to the bead of this barcode. Assuming a simple case of equal number of total human and mouse transcripts in the sample, the cell barcode purity is provided by the formula 1 – impurity where impurity is twice the ratio of transcripts belonging to the minoritary species among the transcripts captured by the cell barcode. Cell barcodes are assigned to each species, mouse or human, if they have a cell barcode purity of 95% or more; for cell barcode purity lower than 95% the cell barcodes are represented as gray dots in the Barnyard plot. The total cell multiplet rate is extrapolated from the measured hetero-species cell multiplet rate (twice the rate in case of an equal number of human and mouse cells), where a hetero-species cell multiplet is defined as a cell barcode for which cell barcode purity is lower than 2/3 (i.e., for which cell impurity is higher than 1/3).

#### B) Post-processing pipeline

The data post-processing pipeline of Cytonaut takes as input the sample count matrix provided by the pre-processing pipeline and performs the following steps:

i. Filtering of cells and/or genes is achieved according to application-dependent parameter values which are selected by the user (e.g., minimum number of cells expressing a gene set to 3, minimum number of genes per cell set to 200, maximum mitochondrial transcript rate per cell set to 20% for the presented studies);
ii. Cell data normalization and log-transformation are applied using the formula log2(10000*X+1)), where X is the percentage of transcripts detected in the cell for the gene of interest and log2() is the logarithm in base 2;
iii. Highly Variable Gene (HVG) detection is performed according to application-dependent parameter values which are decided by the user (e.g., number of top genes set to 2000 for the presented studies);
iv. Principal Component Analysis (PCA) is run on the top HVG genes, according to application-dependent parameter values which are fixed by the user (e.g., number of PCA dimensions set to 50 for the presented studies);
v. 2D Embedding of cells is carried out according to parameter values that are decided by the user (e.g., number of neighbors set to 10 for the presented studies, UMAP method with minimum distance set to 0.3 for the presented studies, or t-SNE method with perplexity set to 30);
vi. Cell Clustering is performed according to parameter values which are set by the user (e.g., Louvain method with resolution set to 0.8 for the presented studies, or Leiden method with resolution set to 0.8);
vii. Differential Gene Expression (DGE), according to a statistical test method established by the user (e.g., Wilcoxon-Mann-Whitney test for the presented studies).

#### C. Post-processing pipeline output

The following items are provided:

i. The DGE matrix, which contains for each cell cluster and each gene the z-score, the log fold change, and the p-values (non-adjusted and adjusted) of the expression of the gene in the cells belonging to the cluster compared to the cells belonging to the other clusters.
ii. The attributes of each cell, including the cell quality indicators and the cluster ID of the cell;
iii. The attributes of each gene, including the cell quality indicators and the gene variability indicators;
iv. The AnnData object (h5ad format), which contains all the data allowing to further reproduce the post-processing results.

#### D) Interactive data visualization

The Cytonaut Rover module provides an interactive visualization of data generated by the post-processing pipeline, allowing exploration of gene expression and cell attributes in cell clusters and application of customized filters leading to customized distribution statistics (heatmap, dot plots, violin plots). In particular, the visualization of the expression of the top differentially expressed genes in each cell cluster facilitates the annotation of the identified cell types.

## Acknowledgements

We gratefully acknowledge helpful discussion with Wilko Duprez, Sahar Sabetnia, Marie-Claude Marchand, Yannick Marie, Laetitia Strehl, Emeline Mundwiller, Justine Guégan, Allison Mallory, Jean-David Bouaziz, and Arturo Londono.

## Author contributions

The first 26 authors all contributed to the development of RevGel-seq and the bioinformatic tools used to analyze the resulting data. The subsequent five authors (N.R, C.B, A.L., G.G, and B.B.) participated in obtaining experimental data from pancreas islet cells. The last two (corresponding) authors (P.W. and S.J.E) directed the research. Drafting of the manuscript was carried out by S.J.E., along with J.K, R.P, A.G., D.D., P.C. and B.A., with participation in the construction of figures by S.L.

## Availability of Data

Sequencing data was deposited on GEO, with the following accession numbers: SuperSeries record GSE218213, with SubSeries records GSE218206,GSE218207,GSE218208,GSE218209,GSE218210,GSE218211.

## Supplementary Information

The online version contains supplementary material available at (link to be added)

---

**Figure S1.**
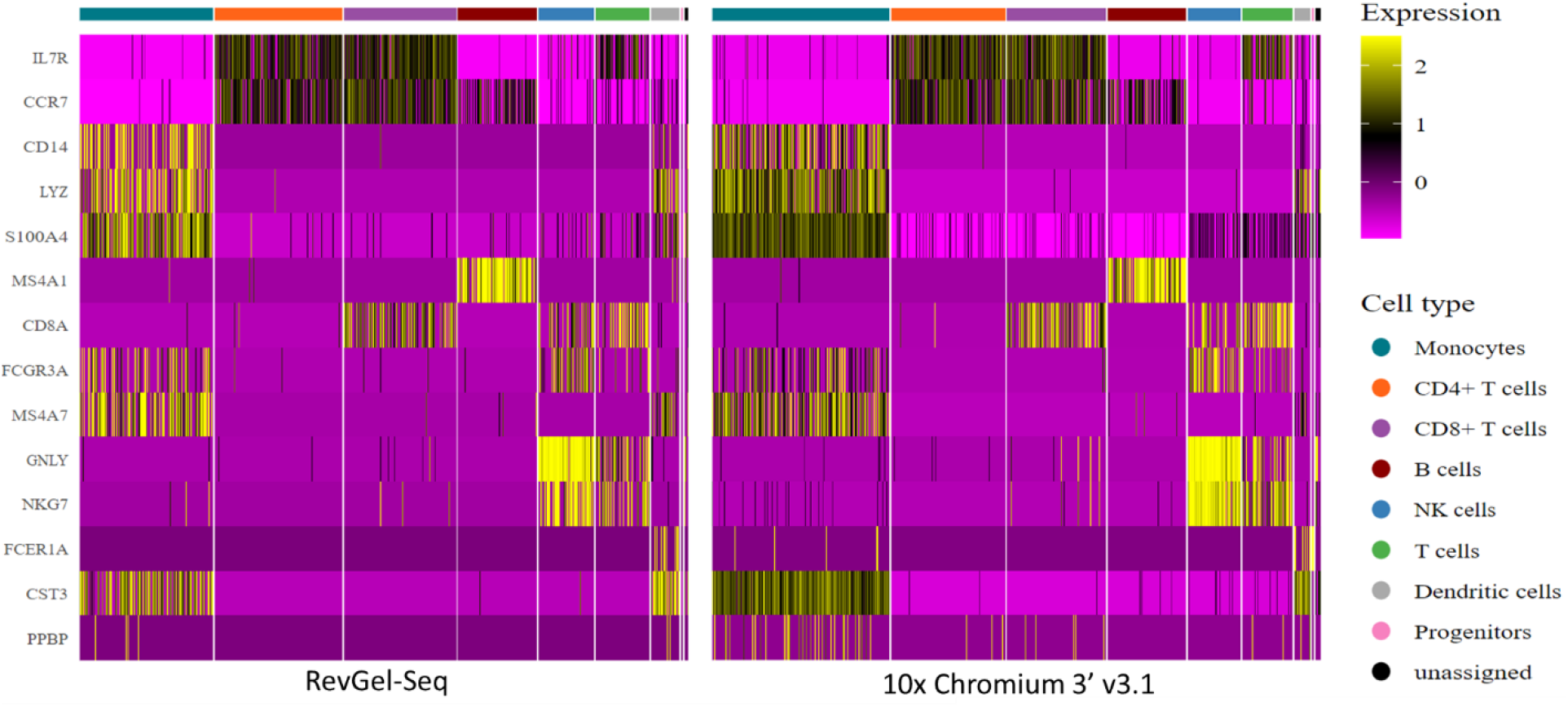
Heatmap of the human PBMC sample prepared with RevGel-seq (left) and with 10x Chromium 3’ v3.1 (right), showing the log-normalized expression of 14 typical canonical PBMC marker genes (shown on the left) in each analyzed cell, where cells are grouped by automatically annotated cell type (8 subtypes, shown on the right) based on the reference dataset MonacoImmuneData. Marker genes: IL7R, CCRT for T cells; CD14, LYZ, S100A4, FCGR3A, MS4A7 for Monocytes; CD8A for CD8+ T cells; MS4A1 for B cells; GNLY, NKG7 for Natural Killer (NK) cells; FCER1A and CST3 for Dendritic cells; PPBP for Platelet cells).

**Figure S2.**
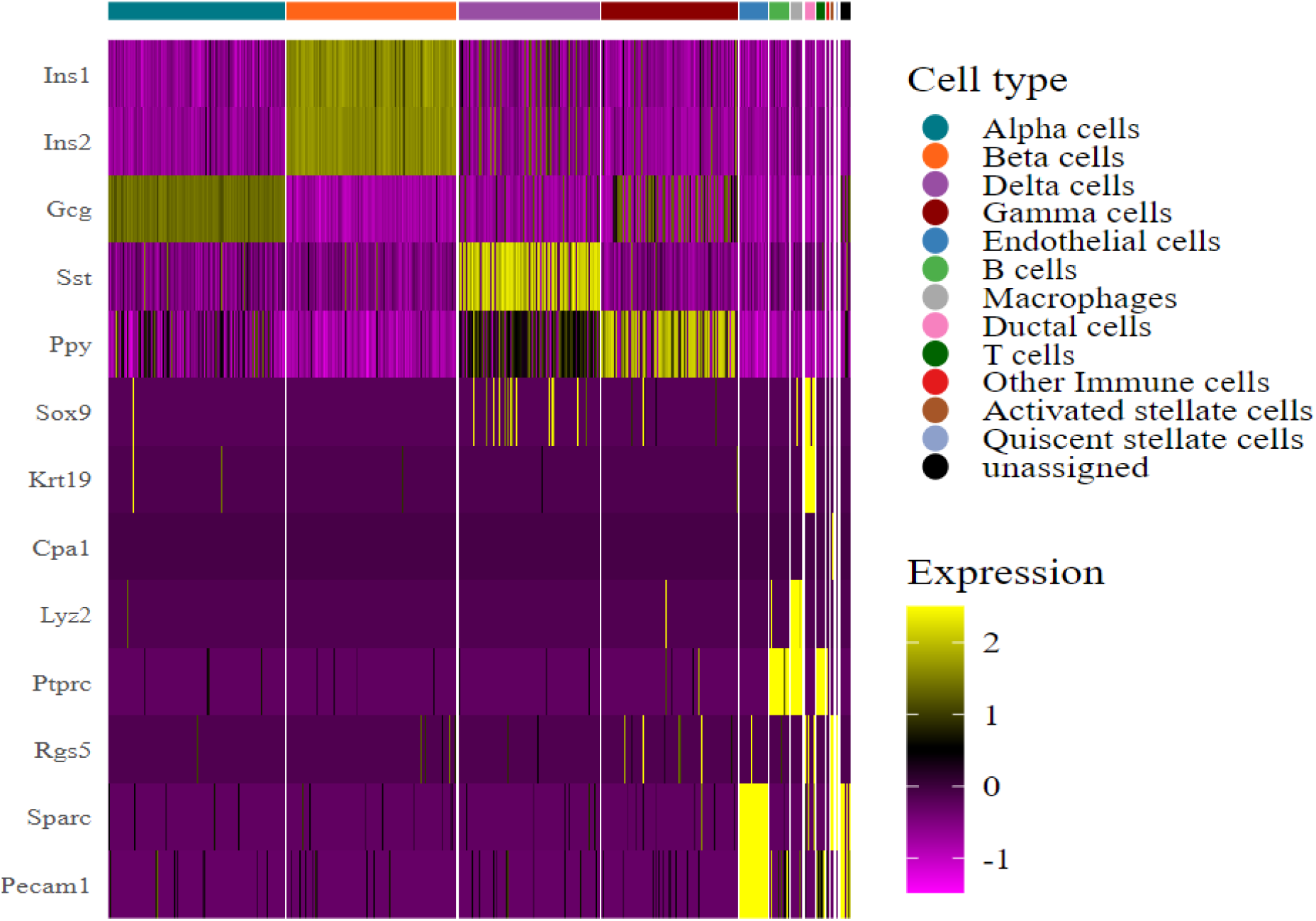
Heatmap of the pancreatic islet cells sample prepared with RevGel-seq, showing the log-normalized expression of 13 marker genes^30^ in each analyzed cell, where cells are grouped by automatically annotated cell type based on the reference dataset BaronPancreasData. Marker genes: Ins1 and Ins2 for Beta cells; Gcg for Alpha cells; Sst for Delta cells; Ppy for Gamma cells; Krt19 for Ductal cells; Cpa1, Lyz2, Ptprc for Acinar cells; RGS5 for Activated stellate cells; Sparc, Pecam1 for Endothelial cells.

**Figure S3.**
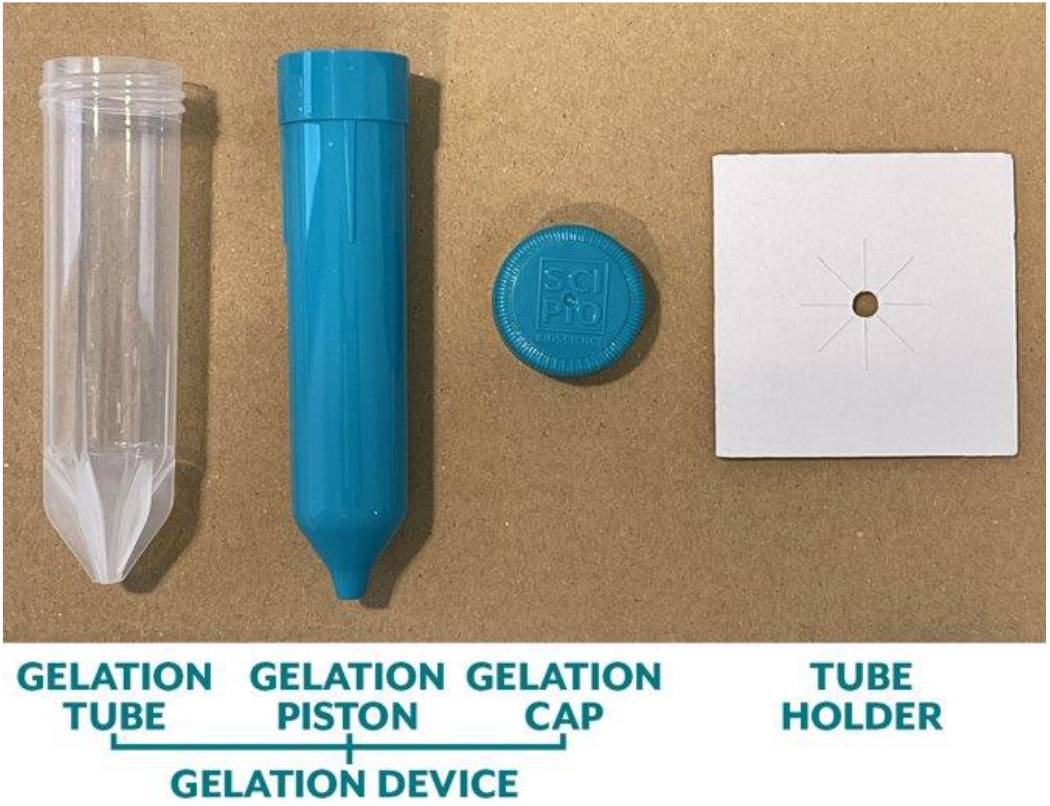
Components of the gelation device: the gelation tube similar in size to a standard 50 mL Falcon conical centrifuge tube, the gelation piston, the gelation cap, and the accompanying tube holder. Spacer guides between the tube and the piston ensure a uniform gel thickness of 500 microns.

**Table S1.**
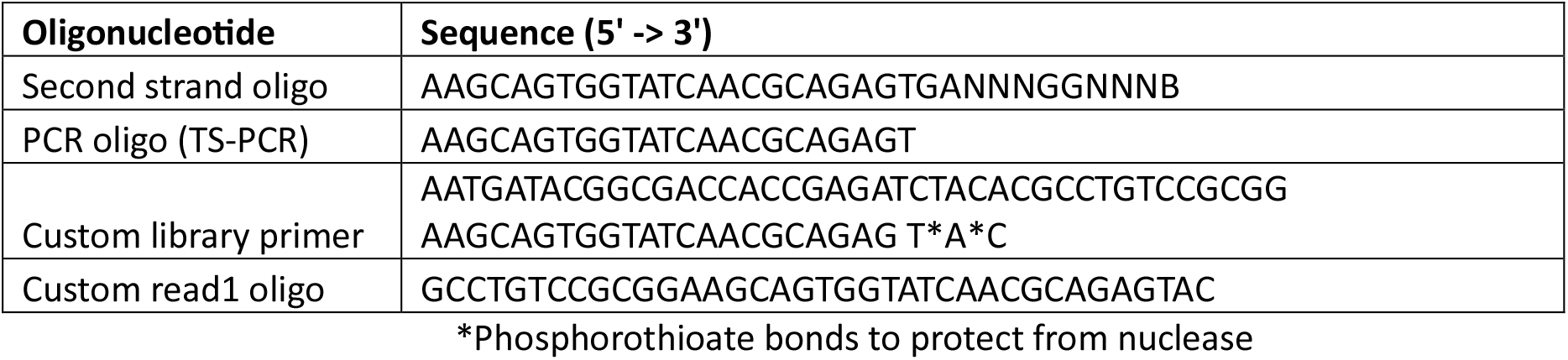
Oligonucleotide sequence for RevGel-seq

**Table S2.** PCR condition for barcoded cDNA amplification

| Step number and description |  | Number of cycles | Temperature | Duration |
| --- | --- | --- | --- | --- |
| Step 1 | Initial denaturation | - | 95°C | 3 min |
| Step 2 | Denaturation | 4 | 98°C | 20 sec |
|  | Annealing |  | 65°C | 45 sec |
|  | Extension |  | 72°C | 3 min |
| Step 3 | Denaturation | 9 | 98°C | 20 sec |
|  | Annealing |  | 67°C | 20 sec |
|  | Extension |  | 72°C | 3 min |
| Step 4 | Final extension | - | 72°C | 5 min |
| Step 5 | Storage | - | 4°C | ∞ |

